# R.ROSETTA: an interpretable machine learning framework

**DOI:** 10.1101/625905

**Authors:** Mateusz Garbulowski, Klev Diamanti, Karolina Smolińska, Nicholas Baltzer, Patricia Stoll, Susanne Bornelöv, Aleksander Øhrn, Lars Feuk, Jan Komorowski

## Abstract

**Motivation:** For machine learning to matter beyond intellectual curiosity, the models developed therefrom must be adopted within the greater scientific community. In this study, we developed an interpretable machine learning framework that allows identification of semantics from various datatypes. Our package can analyze and illuminate co-predictive mechanisms reflecting biological processes.

**Results:** We present R.ROSETTA, an R package for building and analyzing interpretable machine learning models. R.ROSETTA gathers combinatorial statistics via rule-based modelling for accessible and transparent results, well-suited for adoption within the greater scientific community. The package also provides statistics and visualization tools that facilitate minimization of analysis bias and noise. Investigating case-control studies of autism, we showed that our tool provided hypotheses for potential interdependencies among features that discerned phenotype classes. These interdependencies regarded neurodevelopmental and autism-related genes. Although our sample application of R.ROSETTA was used for transcriptomic data analysis, R.ROSETTA works perfectly with any decision-related omics data.

**Availability:** The R.ROSETTA package is freely available at https://github.com/komorowskilab/R.ROSETTA.

**Contact:** (Mateusz Garbulowski), (Jan Komorowski)

## 1 Introduction

Machine learning approaches aim at recognizing patterns and extracting knowledge from complex data. In this work, we aim at supporting the knowledge-based data mining with an interpretable machine learning framework (Doshi-Velez and Kim, 2017; Molnar, 2020). Recently, understanding the complex machine learning classifiers that explain their output is a highly important topic (Azodi, et al., 2020). Here, we developed a framework for interpretable machine learning analysis that is based on rough set theory (Pawlak, 1982). Moreover, we enriched the system with basic statistical measurements that is a unique development with comparison to current state-of-the-art tools. The rough sets methodology has been widely applied to various scientific areas (Jothi, et al., 2019; Kumar and Inbarani, 2018; Zhang, et al., 2014). One of the most important rough sets properties is discovering patterns from complex and imperfect data (Bal, 2013; Bello and Falcon, 2017). In more details, rough set classification theory generates rule-based models that consist of all minimal sets of features that preserve the discernibility. The main representation of this approach is in the form of legible if-then rules that uncover interdependencies among variables also known as features. The rules are often accredited by the two fundamental measures of support and accuracy. The rule support represents the number of samples or objects that satisfy the if-part of the rule, also known as left hand site (LHS). On the other hand, the accuracy represents the fraction of objects from the support set that satisfy the then-part of the rule, also known as right hand side (RHS). In this study, we based the analysis on two algorithms suitable for rule-set estimation. The Johnson reduction method (Johnson, 1974) that is a greedy algorithm and a Genetic reduction method (Wróblewski, 1995) that is based on the theory of genetic algorithms. Estimating the minimal feature subsets (reducts), and later the rules, consists a crucial step towards model transparency. The main advantage of the rule-based modelling is their ability to make explicit their ability to detect co-predictive mechanisms among the features.

Classification models are trained on labeled objects that are *a priori* assigned to them. Here, the universal input structure for machine learning analyses is a decision table (Huysmans, et al., 2011; Kohavi, 1995). A decision table consists of features and objects, that usually represent columns and rows, respectively. Importantly, the last column in a decision table is used as the decision labels. Most of the omics datasets can be represented as a decision table, therefore machine learning analysis can be performed on various case-control studies. In many cases, a feature selection step is necessary prior to the machine learning analysis (Dash and Liu, 1997; Liu and Motoda, 2012). The main goal of feature selection is to reduce the dimensionality of the decision table while simultaneously maintaining the relevant knowledge. Thus, it is recommended to consider feature selection as a standard step prior to the machine learning analysis, especially for big omics datasets.

The ROSETTA software is an implementation for rule-based classification (Øhrn and Komorowski, 1997). It was implemented in C++ as a graphical user interface (GUI) and command line version. ROSETTA has been successfully applied on various studies to model the biomedical datasets (Gil-Herrera, et al., 2011; Komorowski, 2014; Setiawan, et al., 2009). Here, we present a more accessible and flexible implementation of ROSETTA in the form of an R package. R.ROSETTA substantially extends the functionality of the existing software towards analyzing complex and ill-defined bioinformatics datasets. Among others, we have implemented functions such as undersampling, estimation of rule-statistical significance, prediction of classes, retrieval of support sets and various approaches for model visualization (Fig. 1). To the best of our knowledge, there is no framework that allows for such expanded analysis of transparent classification models. To evaluate the performance of R.ROSETTA, we explored rule-based models for a transcriptomic dataset of gene expression measures for autistic and non-autistic subjects.

**Fig. 1.**
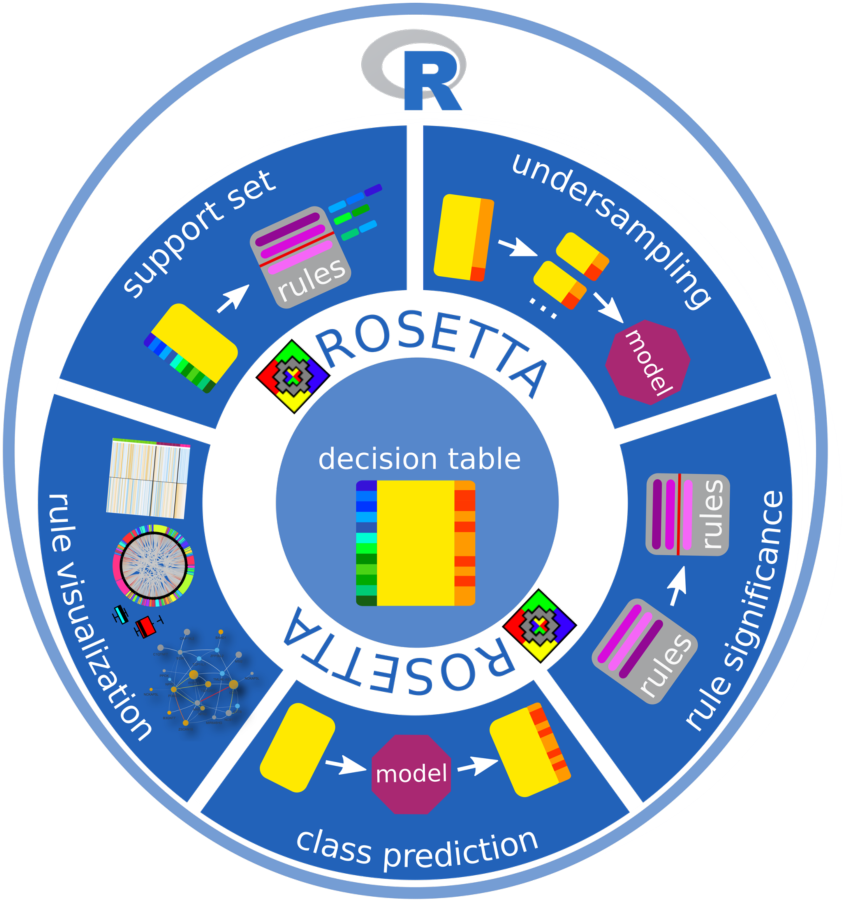
Overview of R.ROSETTA and the major components that were implemented to enhance the ROSETTA functionality.

## 2 Implementation

R.ROSETTA was implemented under R (R Core Team, 2018) version 3.6.0 and the open-source package is available on GitHub (https://github.com/komorowskilab/R.ROSETTA). The R.ROSETTA package is a wrapper (Additional file 1: Package architecture) of the command line version of the ROSETTA system (Øhrn and Komorowski, 1997; Øhrn, et al., 1998). In contrast to ROSETTA, R.ROSETTA is a cross-platform application with multiple additional functionalities (Fig. 1). The succeeding subsections cover a detailed description of the new functions.

### 2.1 Undersampling

Overrepresentation of a decision class over others may lead to biased performance of the model (Additional file 1: Fig. S1). Ideally, each decision class shall contain approximately the same number of objects. To tackle this, we suggested to randomly sample a sufficient number of times the majority class without replacement in order to achieve an equal representation of classes. This approach of balancing the data is generally known as undersampling (Liu, et al., 2009).

To build a balanced rule-based model, we have implemented an option that divides the dataset into subsets of equal sizes by undersampling the larger sets. By default, we require each object to be selected at least once, although the user can specify a custom number of sampled sets, as well as a custom size for each set. Classification models for each undersampled set are merged into a single model that consists of unique rules from each classifier. The overall accuracy of the model is estimated as the average value of the sub-models. Finally, the statistics of the merged rule-set shall be recalculated on the original training set using the function *recalculateRules*.

### 2.2 Rule significance estimation

The p-value is a standard measure of statistical significance in biomedical studies. Here, we introduced p-value estimation as an additional quality measure for the rules. Classification models generated by R.ROSETTA consist of sets of varying number of rules estimated by different algorithms. In the case of the Johnson algorithm, this set contains a manageable number of rules (Table 1, Additional file 1: Table S1), while in the case of the Genetic algorithm this set can be considerably larger (Additional file 1: Table S1, Table S2). In both cases, supervised pruning of rules from the models would not heavily affect the overall performance of the classifier. To better assess the quality of each rule we assume a hypergeometric distribution to compute p-values (Hvidsten, et al., 2005) followed by multiple testing correction. The hypergeometric distribution estimates the representation of the rule support against the total number of objects. When estimating the p-value for a rule, the hypergeometric distribution is adapted to the rule concepts:

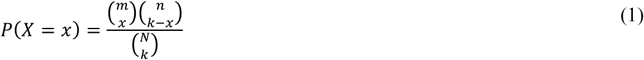

where *x* is the RHS support of the rule, *k* is the LHS support of the rule, *m* is the number of objects matching the decision class for a given rule, *n* is the number of objects for the decision class(es) opposite to a given rule, and *N* is the total number of objects. Such optimized models illustrate the essential co-predictive mechanisms among the features and can be pruned based on their significance levels. We also implemented additional model-tuning statistical metrics for rules including risk ratio, risk ratio p-value and risk ratio confidence intervals (Nakazawa and Nakazawa, 2019). The complete statistical measurements are included in the output of the R.ROSETTA model.

**Table 1.** Performance evaluation for the Johnson reduction method. The average statistic values of support, accuracy and coverage are presented in the table. Top ranked co-predictors were selected as the most significant by Bonferroni-adjusted p-value.

| class | control |  | autism |  |
| --- | --- | --- | --- | --- |
| total number of rules | 207 |  | 194 |  |
| rules | basic | recalculated | basic | recalculated |
| number of rules<br>*($P \leq 0.05$) | 150 | 89 | 128 | 94 |
| LHS support | 13 | 18 | 13 | 16 |
| RHS support | 13 | 17 | 13 | 14 |
| accuracy | 0.97 | 0.94 | 0.98 | 0.85 |
| LHS coverage | 11% | 22% | 11% | 25% |
| RHS coverage | 22% | 20% | 22% | 21% |
| top ranked co-predictors | PPOX, NCS1 | MAP7, NCKAP5L | RHPN1, ZFP36L2 | NCS1, CSTB |

### 2.3 Vote normalization in the class prediction

Rule-based models allow straightforward class prediction of unseen data using voting. Every object from the provided dataset is fed into the pre-trained machine-learning model and the number of rules for which their LHS is satisfied are counted in. In the final step, the votes from all rules are collected for each individual object. Typically, an object is assigned to the class with the majority of votes. However, for some models an imbalanced number of rules for each decision class over another may have been generated. For such cases, we propose adjusting for the rule-imbalance by normalizing the vote counts. We implemented various vote normalization methods in R.ROSETTA. Vote normalization can be performed by dividing the number of votes by its mean, median, maximum, total number of rules or square root of the sum of squares. We compared the performance of these methods in (Additional file 1: Table S3).

### 2.4 Rule-based model visualization

The model transparency is an essential feature that allows visualization of co-predictive mechanisms in local (single rule) and global (whole model) scale. The package provides several ways for visualizing single rules, including boxplots and heatmaps (Additional file 1: Fig. S2D, Fig. S3) that illustrate the continuous levels of each feature of the selected rule for each object. Such rule-oriented visualizations gather the objects into those that belong to the support set for the given class, those that do not belong to the support set for the given class and the remaining objects for the other classes. Such graphic representations can assist towards the interpretation of individual rules of interest and visualization of interactions with respect to their continuous values.

A more holistic approach displays the entire model as an interaction network (Shmulevich, et al., 2002). The R.ROSETTA package allows exporting the rules in a specific format which is suitable with rule visualization software such as Ciruvis (Bornelöv, et al., 2014) or VisuNet (Additional file 1: Fig. S2B) (Anyango, 2016). Such model can be pruned to display only the most relevant co-predictive features and their levels. These approaches provide a different point of view on the interpretation of rule-based models that allow discovering known proof-of-concept and novel interdependencies among features (Dramiński, et al., 2016; Enroth, et al., 2012).

### 2.5 Recapture of support sets

R.ROSETTA is able to retrieve support sets that represent the contribution of objects to rules (Additional file 1: Fig. S2D, Fig. S3). As a result, each rule is characterized by a set of objects that fulfill the LHS or RHS support. For example, in case of the gene expression data, gene co-predictors will be represented with the list of corresponding samples (patients). There are several advantages into knowing this information for the corresponding objects. Support sets contribute to uncovering objects whose levels of features might have shared patterns. Such sets may be further investigated to uncover specific subsets within decision classes. Moreover, non-significant support sets allow detecting objects that may potentially introduce a bias to the model and might be excluded from the analysis.

### 2.6 Synthetic data

To evaluate rule-based modelling with R.ROSETTA, we implemented a function to create synthetic data. The synthetic dataset can be generated with a predefined number of features, number of objects and proportion of classes. Additionally, the user may choose between continuous and discrete data. The synthetic data structure is formulated as a decision table that follows the description in the introduction. A synthetic dataset is constructed from the transformation of randomly generated features computed from a normal distribution. In this approach, the randomly generated features are multiplied by the Cholesky decomposition of positive-definite covariance matrix (Pourahmadi, 1999). The Cholesky decomposition *D* of the matrix *L* is calculated as:

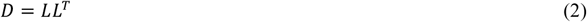

where *L* is a lower triangular covariance matrix, and *LT* is conjugate transpose of *L*.

## 3 Results and discussion

### 3.1 Benchmarking

We evaluated R.ROSETTA against other R packages that perform rule-based machine learning including C50 (Kuhn, et al., 2018), RoughSets (Riza, et al., 2014) and RWeka (Hornik, et al., 2009) (Additional file 1: Benchmarking, Table S4). We compared the efficacy of the classification algorithms by measuring the accuracy, the ROC area under curve (AUC), the running time and the total number of rules (Fig. 2, Additional file 1: Fig. S4). To perform compatible benchmarking across algorithms, we standardized the classification procedure for equal frequency discretization and 10-fold cross validation (CV). Additionally, to account for the stochasticity introduced by sampling in CV, each algorithm was executed 20 times with different seed values.

**Fig. 2.**
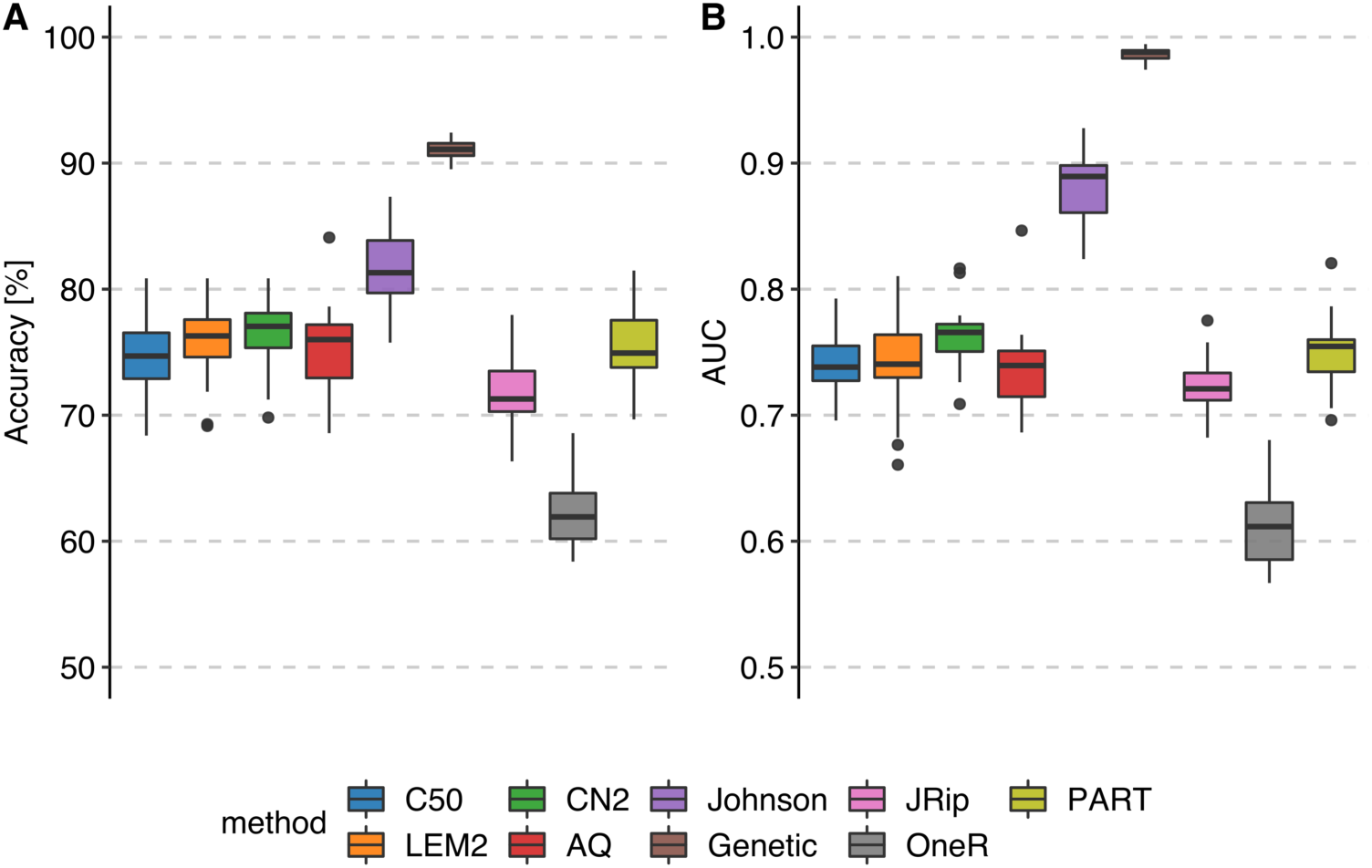
Benchmarking the 9 rule-based machine learning methods. **A** Average accuracy of the models. **B** Average ROC AUC of the models.

Even though R.ROSETTA produced the highest quality models (Fig. 2), its runtime, especially for the genetic algorithm, was higher than other algorithms (Additional file 1: Fig. S4B). However, the runtime of computations strongly depends on the algorithm complexity. Most of the systems compute one reduct or one set of features which is of linear complexity while R.ROSETTA computes all reducts, which is an NP-hard problem. We believe this is an important feature since living systems are robust and usually have alternative ways of achieving the outcome. Clearly, as a consequence of estimating multiple reducts, R.ROSETTA algorithms tend to produce more rules in comparison to other methods (Additional file 1: Fig. S4A). It is then natural to remove the weakest rules. To this end, we suggest to prune the rule-set using the quality measurements such as, for instance, support or p-value.

We observed that the surveyed packages do not provide quality-statistic metrics for the model and the rules, in particular. In contrast, the R.ROSETTA package includes a variety of quality and statistical indicators for models (accuracy, AUC) and rules (support, p-value, risk ratio etc.) in an effortlessly and R-friendly inspectable output. Furthermore, the evaluated packages do not include novel R.ROSETTA features such as undersampling, support sets retrieval and rule-based model visualizations.

### 3.2 Sample application

To demonstrate R.ROSETTA in a bioinformatics setup, we focused on transcriptomic data analysis. We examined gene expression levels of 82 autistic and 64 non-autistic (control) male children (Additional file 1: Table S5) downloaded from the GEO repository (GSE25507) (Alter, et al., 2011). The expression of 54678 genes was measured with the Affymetrix Human Genome U133 Plus 2.0 array. The dataset was preprocessed (Additional file 1: Data preprocessing) and corrected for the effect of age (Additional file 1: Fig. S5). The decision table was ill-defined with the number of features (in this case genes) being much larger than the number of samples. To handle the high dimensionality, we employed the Fast Correlation-Based Filter method (Novoselova, et al., 2018) that performs feature selection (Additional file 1: Feature selection, Table S6). The final decision table was reduced to 35 genes (Additional file 1: Table S7).

We constructed two balanced models (Additional file 1: Classification) with R.ROSETTA for Johnson and Genetic reducers with 82% and 90% accuracy (0.85 and 0.98 area under the ROC curve), respectively (Fig. 3A, Additional file 1: Fig. S2A, Table S1). Even though the overall performance of the Genetic algorithm was better than Johnson’s, its tendency to generate numerous rules reduced the significance of individual rules after correcting for multiple testing (Fig. 3C, Additional file 1: Fig. S6, Table S1).

**Fig. 3.**
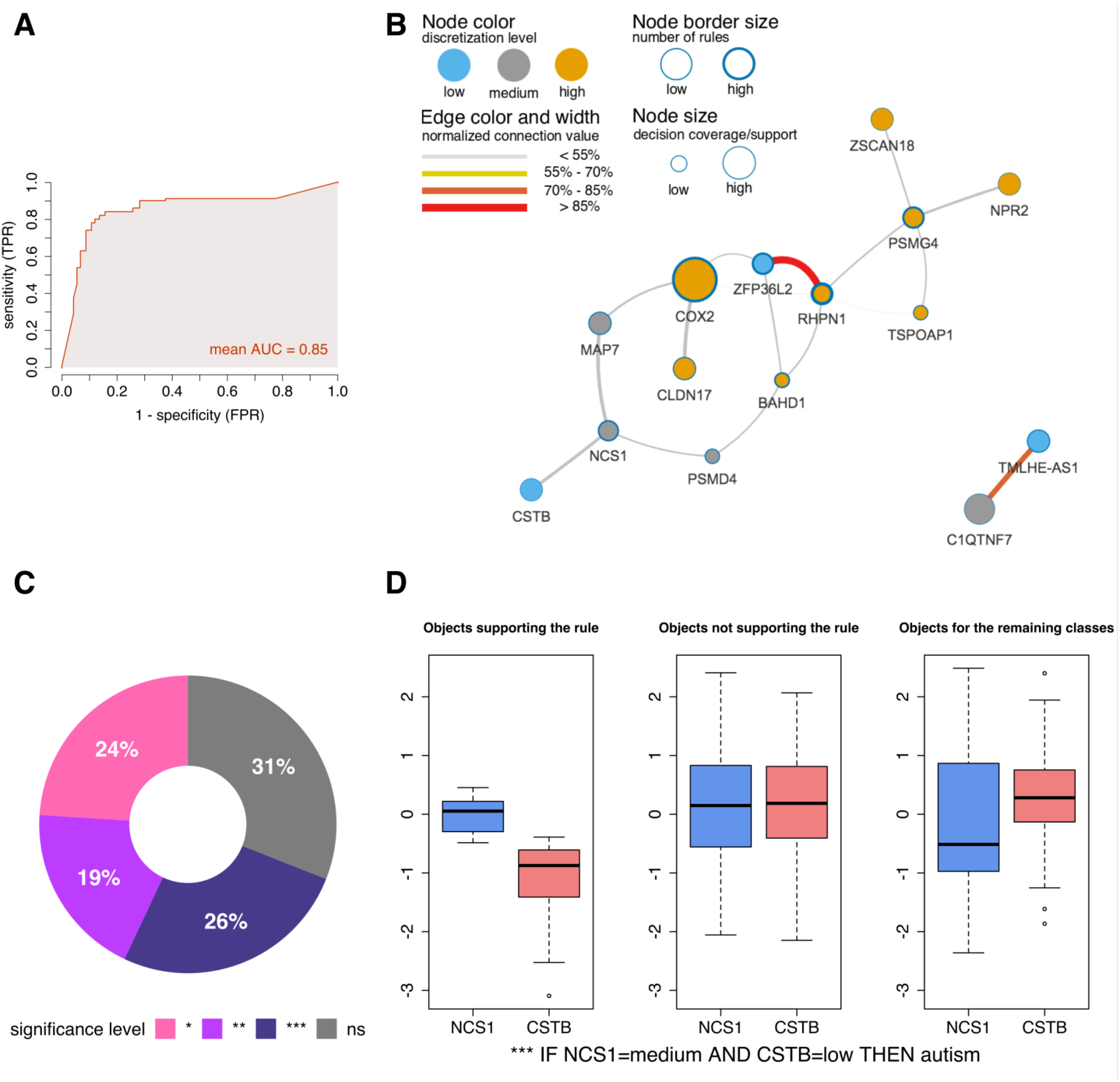
Rule-based model for autism-control performed with the Johnson reduction method **A** ROC AUC) for the model. **B** VisuNet network of feature interdependencies for the autism class. Rules were selected based on their statistical significance (Bonferroni-adjusted P ≤ 0.05). **C** Distribution of the significance of rules in the model. Bonferroni-adjusted p-values are marked with ns(P > 0.05), *(P ≤ 0.05), **(P ≤ 0.01) and ***(P ≤ 0.001). **D** Distribution of support sets for the top-ranked rule from the recalculated model. Two left-most boxplots show objects supporting and non-supporting the autism class. The right-most boxplot shows objects from the control class.

To identify the most relevant interdependencies among genes, we selected significant rules (Bonferroni-adjusted P ≤ 0.05) (Fig. 3D, Table 1) from the Johnson model. The highest ranked co-predictors include the medium expression levels of neuronal calcium sensor 1 (*NCS1*) and low expression levels of cystatin B (*CSTB*). The *NCS1* gene is related to the calcium homeostasis control (Boeckel and Ehrlich, 2018). Dysregulated expression and mutations in *NCS1* have been linked to neuropsychiatric disorders (Boeckel and Ehrlich, 2018). Moreover, previous studies have demonstrated that calcium homeostasis is altered in autism disorders (Palmieri, et al., 2010), while the second feature of the rule, the reduced expression of *CSTB* has been linked to the mechanism of pathogenesis in epilepsy (Lalioti, et al., 1997). This co-predictive mechanism is additionally characterized by negative Pearson’s rank correlation coefficient (Additional file 1: Fig. S7).

We also utilized the VisuNet framework that supports visualization and exploration of rule-based models. Moreover, we displayed a rule-based network for the significant rules for autism (Bonferroni-adjusted P ≤ 0.05) (Fig. 3B). The biggest node in the network is the cyclooxygenase 2 (*COX2*) gene and suggests a meaningful contribution to the prediction of autistic young males. Elevated expression of *COX2* has been earlier associated with autism (Yoo, et al., 2008). The study reported that *COX2* carried the Single Nucleotide Polymorphism (SNP) rs2745557 and the GAAA haplotype that were significantly associated with autism (Yoo, et al., 2008). Moreover, *COX2* is constitutively expressed in neuronal tissues of patients with psychiatric disorders (Ibuki, et al., 2003). Based on the network, we can also notice a very strong interdependency between the high expression levels of rhophilin rho GTPase binding protein 1 (*RHPN1*) and the low expression levels of *ZFP36* ring finger protein like 2 (*ZFP36L2*). The association of abnormalities in the GTPase signaling pathway and neurodevelopmental disorders has been previously reported (Reichova, et al., 2018). Rho GTPases participate in the spectrum of signaling pathways related to neurodevelopment such as neurite extension or axon growth and regeneration (Reichova, et al., 2018). The second component is a zinc-finger protein coding gene. The enrichment of lowly expressed zinc fingers in the case-control studies of autism was also discovered by the authors of this dataset (Alter, et al., 2011). We investigated additional autism-related genes that have been reported in the supplementary material (Additional file 1: Feature validation). The described above co-prediction mechanisms illustrate links among the autism-related genes that may explain genetic interactions. Moreover, we found co-prediction mechanisms among the genes that were not previously reported as autism-related genes e.g. proteasome assembly chaperone 4 (*PSMG4*) or bromo adjacent homology domain containing 1 (*BAHD1*) (Fig. 3B). Although the above analysis was performed for transcriptomic data, we have shown that ROSETTA and R.ROSETTA approach performs very well for various other omics data types e.g. DNA methylation (Moghadam, et al., 2016).

### 3.3 Synthetic data evaluation

To explore the influence of the basic concepts of the decision table into the rule-based modelling, we implemented a function that generates synthetic decision tables. We used the synthetic data to describe the rule-based model performance with regards to the number of features, the number of objects and the decision-class imbalance (Additional file 1: Fig. S8). Increasing the number of features did not affect the quality of the model that remained stable across tests (Additional file 1: Fig. S8B, Fig. S8C). However, increasing the number of objects moderately improved the overall quality of the model (Additional file 1: Fig. S8E, Fig. S8F). Furthermore, we showed that undersampling corrected the bias that arose from the class imbalance (Additional file 1: Fig. S1). We found that in case of modelling data with class imbalance, it is better to compare the AUC measure rather than the accuracy (Additional file 1: Fig. S1). Overall, tests on synthetic data showed the performance of the rule-based models under different settings. This will help improve the understanding of rule-based machine learning modelling.

## 4 Conclusions

In this work, we developed R.ROSETTA, a framework that assists analysis of interpretable machine learning models. R.ROSETTA consists of fundamental components of statistics for rule-based modelling and allows for more accessible machine learning studies. The package significantly enhances the accessibility for R users to the transparent machine learning environment and the interpretability of the results. Furthermore, the original ROSETTA functions have been improved and adapted to the R programming environment by an incorporation of novel components targeting bioinformatics applications. These improvements include undersampling datasets to account for imbalances, estimation of the statistical significance of classification rules, retrieving objects from support sets, prediction of labels of external datasets and integration with rule-based visualization tools. We assessed the performance of R.ROSETTA on a complex dataset involving gene expression measurements for autistic and non-autistic young males. We demonstrated that R.ROSETTA facilitated the detection of novel interdependencies for autism-related genes. Additionally, by testing the package with synthetic data, we described basic concepts of rule-based learning. The results demonstrated the potential of transparent machine learning modelling that can be performed with R.ROSETTA.

## Acknowledgments

We would like to thank F. Barrenäs, S. Belin, M. Cavalli, Z. Khaliq, B. T. Moghadam, G. Pan, C. Wadelius and S. Younes for their insightful discussions and testing/debugging the package.

