## Supplementary Material for "R.ROSETTA: an interpretable machine learning framework"

### R.ROSETTA: a framework for interpretable machine learning modelling

[1]Department of Cell and Molecular Biology, Uppsala University, Sweden, [2]Department of Immunology, Genetics and Pathology, Uppsala University, Sweden, [3]Department of Research, Cancer Registry of Norway, Norway, [4]Department of Biosystems Science and Engineering, ETH Zurich, Switzerland, [5]Cancer Research UK Cambridge Institute, University of Cambridge, UK, [6]Department of Informatics, University of Oslo, Norway

*To whom correspondence should be addressed

#These authors contributed equally to the work as second authors.

 (Mateusz Garbulowski)

 (Jan Komorowski)

R.ROSETTA is freely available at: https://github.com/komorowskilab/R.ROSETTA

Tutorials and more information can be found at: https://komorowskilab.github.io/R.ROSETTA/

### 1. Notes

#### 1.1. Package architecture

The ROSETTA framework comes in a GUI version for Windows systems and a command line version for UNIX-based systems. The R.ROSETTA package is a cross-platform application that uses command line ROSETTA. However, UNIX-based systems require installation of the compatibility layer software Wine (https://www.winehq.org/). For more information we recommend to read the original ROSETTA articles (Øhrn, 1999; Øhrn, 2000; Øhrn, et al., 1998) and the technical reference manual (Øhrn, 2001). Detailed instructions for the R.ROSETTA installation, functions and a sample code are available in the package manual or on the official R.ROSETTA website (https://komorowskilab.github.io/R.ROSETTA/).

#### 1.2. Benchmarking

We compared R.ROSETTA with three other R packages designed for rule-based modelling. In total, we compared 9 different algorithms. We excluded algorithms that focus on fuzzy rule-based learning, operate on continuous decision classes and/or include fixed internal discretization methods. We standardized the procedure of running the algorithms so that each method was performed with 10-fold CV and equal frequency discretization. The parameters of each methods were set to default. Each method was repeated 20 times with a different seed level. The runtime of each method was measured as a time required for inputting a data to the function to generating a rule-based model. In the discretization part, the cuts from a training part were applied to discretize the test set. The final values for all the measures were calculated as an average value from 20 repetitions. The calculations were performed with macOS High Sierra with the following parameters: processor 2,2 GHz Intel Core i7 and memory 8GB 1600MHz DDR3.

#### 1.3. Data preprocessing

The so-called autism-control dataset was loaded and processed using the getGEO function from the GEOquery R library (Davis and Meltzer, 2007). The data was normalized with the Robust Multi-array Average (RMA) functions *ReadAffy* and *rma* from the affy R package (Gautier, et al., 2004). Gene names were retrieved using AnnotationDbi (Pages, et al., 2008) and hgu133plus2.db (Carlson, 2016) R packages. To identify unknown probe names, the annotation table HG-U133_Plus_2.na36 downloaded from http://www.affymetrix.com/site/mainPage.affx was processed and the probe coordinates were intersected with the unknown probes using the GenomicRanges (Lawrence, et al., 2013) R package. For one probe the gene name could not be identified due to the lack of coordinates. The clinical data was investigated for potential batch effects using the Pearson correlation with the rcorr function from the Hmisc R library (Harrell and Dupont, 2008). The age of the samples was highly correlated to the outcome. The sva R package (Leek, et al., 2012) was used to correct the data for the age effect.

#### 1.4. Feature selection

Dimensionality reduction of the autism-control dataset was performed with the Fast Correlation-Based Filter (FCBF) method (Yu and Liu, 2003) available as a function in the Biocomb R package (Novoselova, et al., 2018). The feature selection method utilized the predominant correlation of the features and decision along with the redundancy. We chose FCBF as the method that selected the highest number of important features among the tested software. The other advantage of using FCBF was the compatibility of the discretization method with R.ROSETTA. The FCBF function was set to Equal Frequency discretization for 3 states. The method selected 35 features with Information Gain above 0. This step could be alternatively performed with other dimensionality reduction methods such as Monte Carlo Feature Selection (Dramiński, et al., 2010; Dramiński and Koronacki, 2018), Boruta (Kursa and Rudnicki, 2010), Student's t-test, caret R package (Kuhn, 2008) and many others.

#### 1.5. Classification

The final decision table was constructed using the 35 important genes, 146 objects/samples and the decision class (autism or control). The models were created using 10-fold cross validation (CV) for the standard voter method, equal frequency discretization into 3 states, discernibility of the objects and Bonferroni method to adjust rule p-values for multiple testing. Undersampling was applied to remove a slight imbalance between the decision classes. The paper presents Johnson and Genetic reduction methods. However, for simplicity of the model and rule significance reasons most of the interpretations were based on the rules estimated from the Johnson reducer.

#### 1.6. Feature validation

We described sample genes that were likely associated with autism in the results section of the main article. Additionally, we depicted genes that had been earlier linked to the brain or the nervous system such as: migraine, headache – PPOX (Makki, et al., 2017), smell perception – OR51B5 (Chen, et al., 2018) and ataxia – ATXN8OS (Moseley, et al., 2006). We found that NCKAP5L is a gene likely to be involved in neurodevelopmental dysfunction in autism (Chahrour, et al., 2012). We showed that expression of NCKAP5L gene was down-regulated in non-autistic patients. Among the most relevant features, we discovered a group of zinc fingers (Alter, et al., 2011) such as ZSCAN18/ZNF447, ZFP36L2, and KLF8/ZNF741 and a group of genes related to calcium homeostasis control (Palmieri, et al., 2010) such as SCIN, NCS1, and CAPS2.

#### 2. Figures

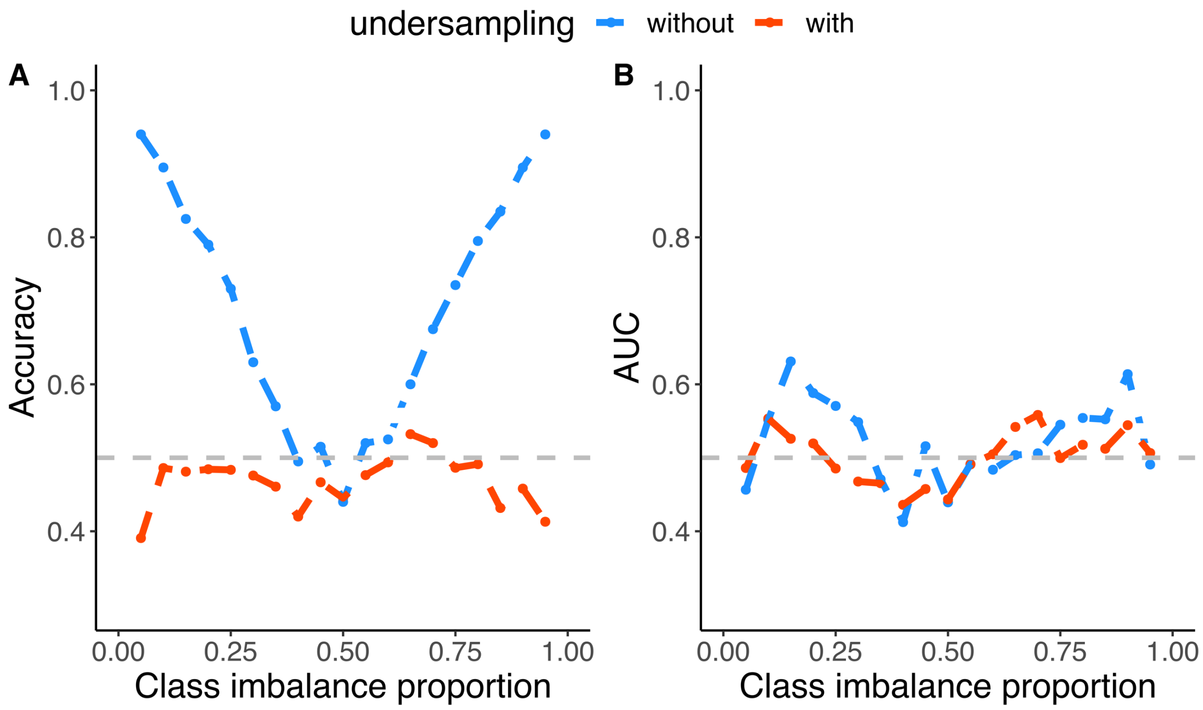

**Fig. S1.** An undersampling application for various class imbalance proportions. The tests were performed with the Johnson reduction algorithm on the synthetic data with a random correlation (expected accuracy 50%). The synthetic data consisted of 50 features and 100 objects. The class imbalance proportion was established from 0.05 to 0.95 with a 0.05 step. **A** The relation between the class imbalance and model accuracy **B** The relation between the class imbalance and the area under the ROC curve.

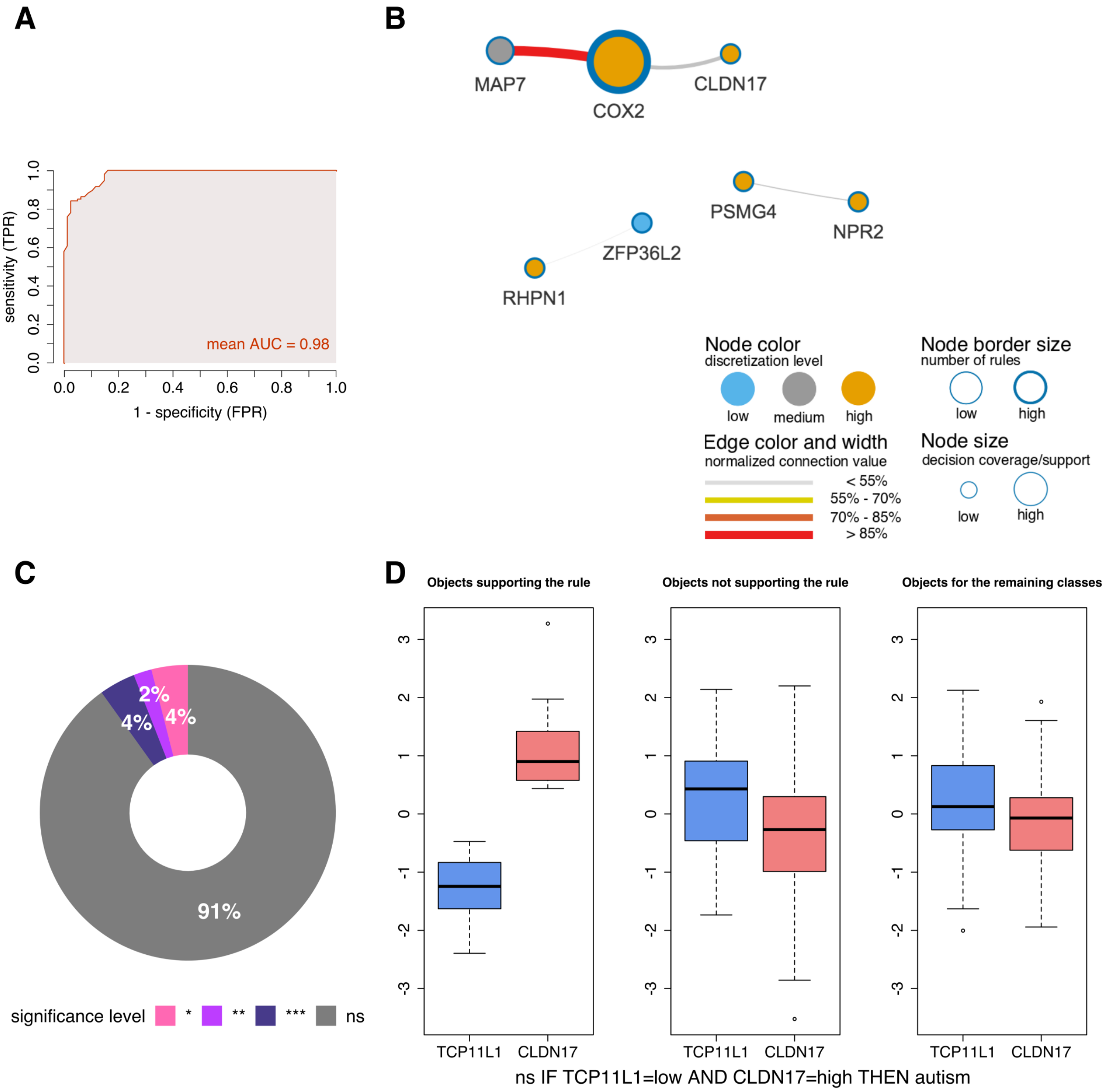

**Fig. S2.** Rule-based model for the autism-control performed with the Genetic reduction method **A** Receiver operating characteristic curve (ROC) and area under the ROC curve (AUC) value of the model. **B** VisuNet network of feature interdependencies for the autism class. The rules were selected according to $P \leq 0.05$. **C** Distribution of the rule significance levels in the model. Bonferroni-adjusted rule p-value is marked with ns($P > 0.05$), *($P \leq 0.05$), **($P \leq 0.01$) and ***($P \leq 0.001$). **D** Distribution of support sets for the top ranked rule from the recalculated model. Two left-most boxplots present objects supporting and non-supporting the autism class. The right-most boxplot presents objects from control class.

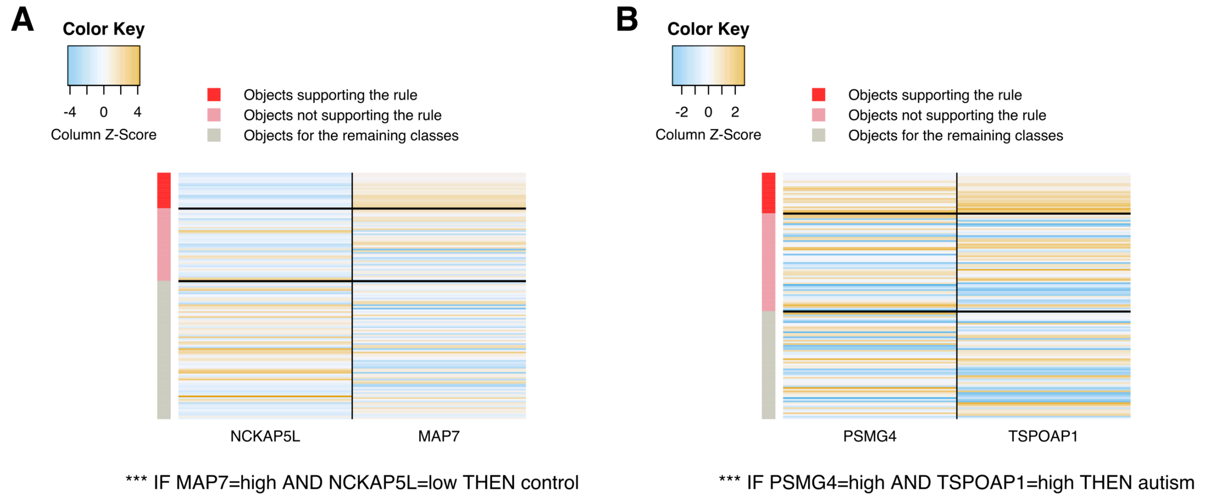

**Fig. S3.** A rule-oriented graphic representation of its corresponding continuous values from the decision table. A given rule comes from the recalculated autism-control model. **A** The most significant co-predictors for the control class **B** The most significant co-predictors for the autism class

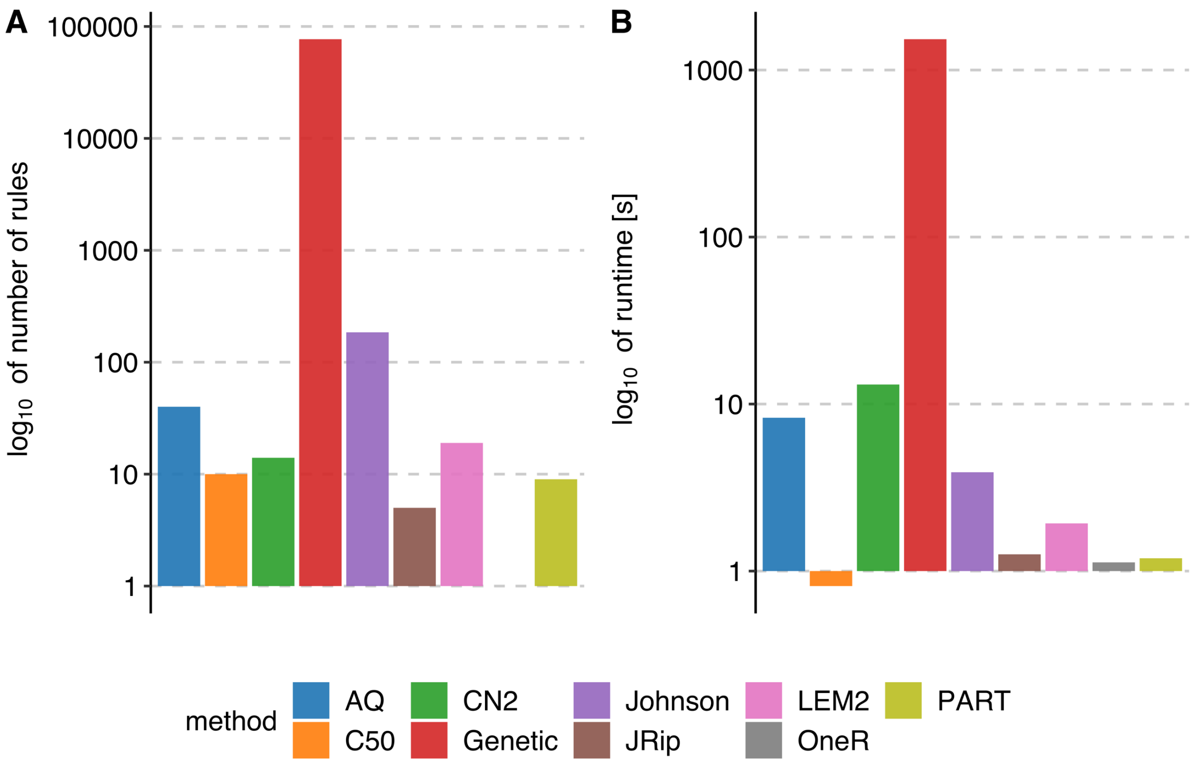

**Fig. S4.** A comparison of the rule number and runtime of nine rule-based machine learning methods. A The number of rules in the model. B The runtime of the algorithm, measured from the time required for inputting a decision table to receiving a rule-based model.

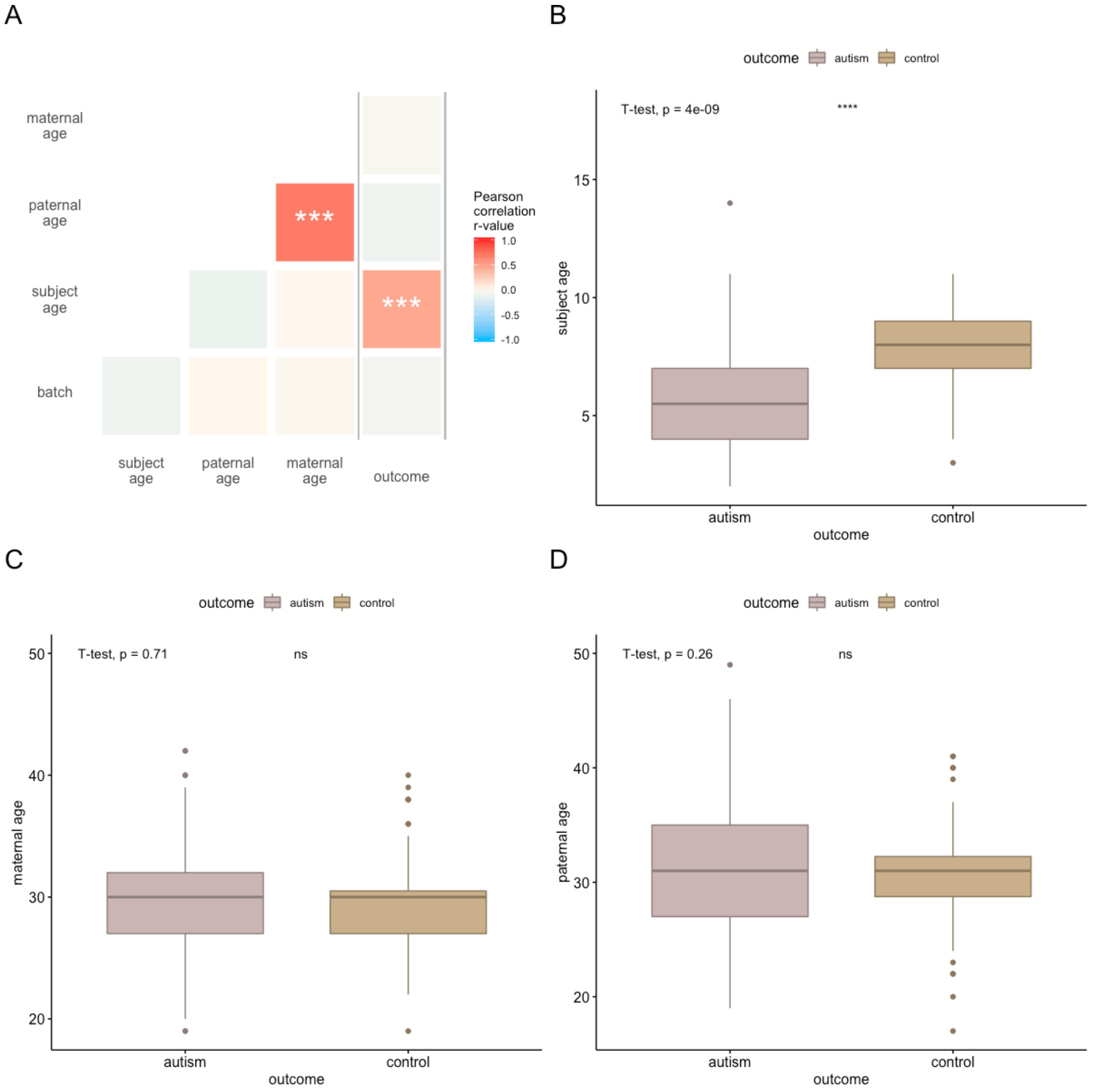

**Fig. S5. A** Pearson correlation r values derived from the clinical data. Stars represent the p-value significance level of the correlation values, denoted by non-significant: ns($p > 0.05$) and significant: *($p \leq 0.05$), **($p \leq 0.01$), ***($p \leq 0.001$). The decision column is detached with dark blue lines to illustrate the effect of the clinical data on the decision. **B** Relationship between outcome and subject age **C** Relationship between the outcome and maternal age. **D** A relationship between outcome and paternal age. The Student's t-test was used to test for a difference in subject, maternal or paternal age between cases and controls in **B**, **C** and **D** with p-values indicated in the figure.

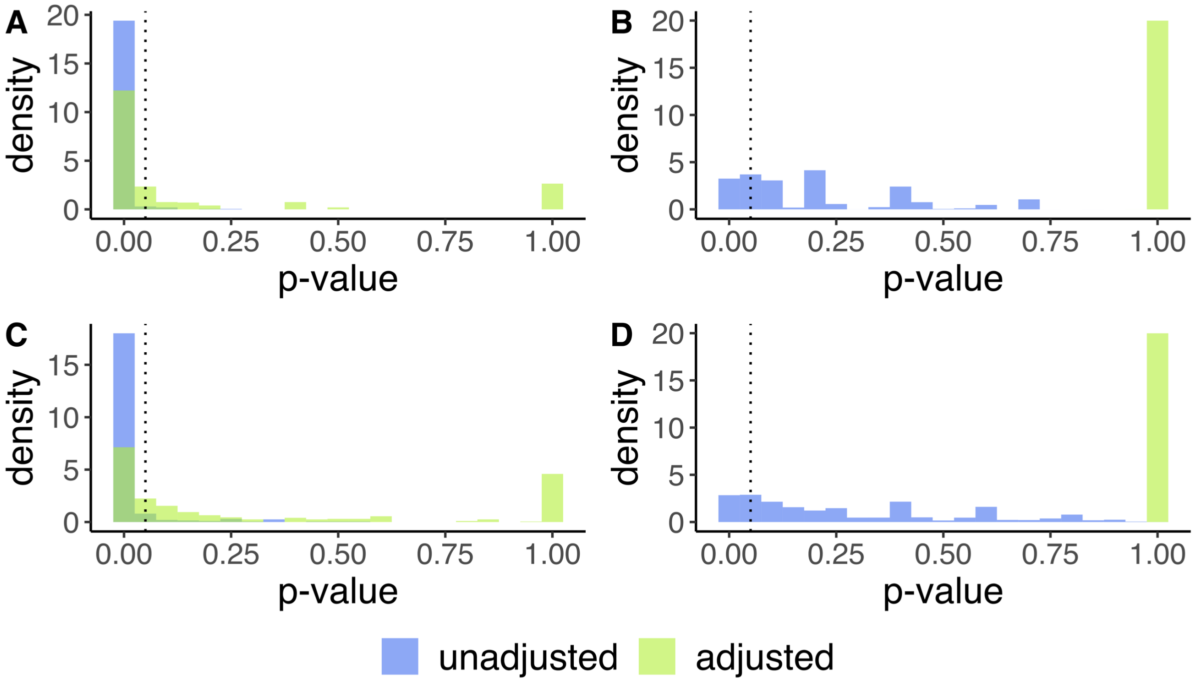

**Fig. S6.** The density of rule p-values for the reduction methods. Histograms display the comparison of the p-value adjustment and the model recalculation between reducers. A dotted line marks the 0.05 significance threshold. **A** autism-control basic model generated with the Johnson reducer method **B** autism-control basic model generated with the Genetic reducer method **C** autism-control recalculated model with the Johnson reducer method **D** autism-control recalculated model with the Genetic reducer method.

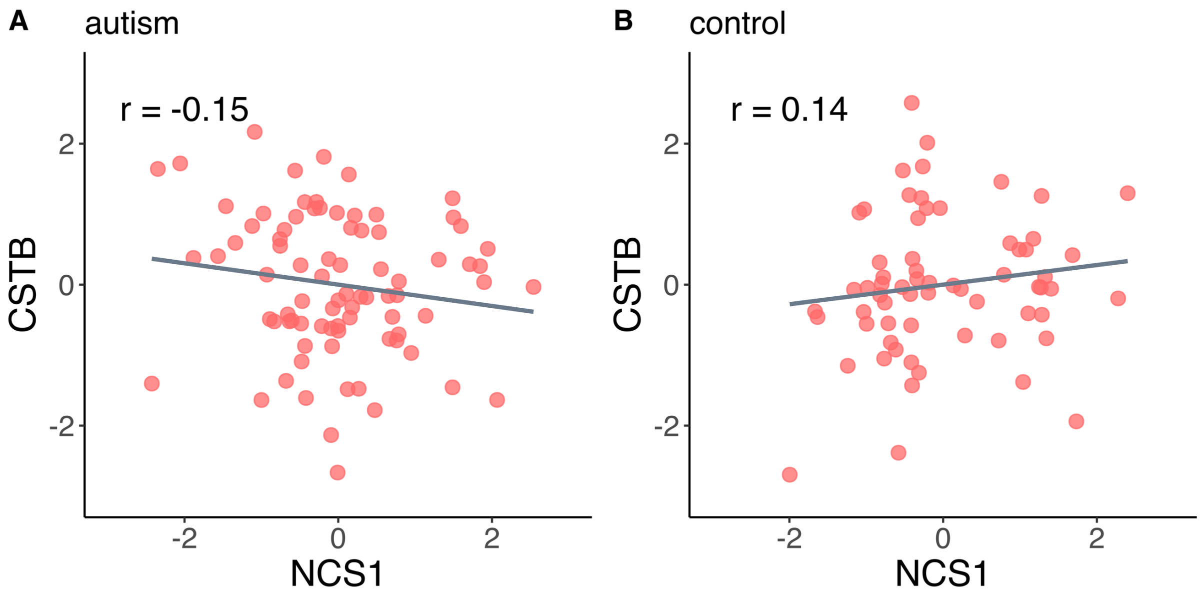

**Fig. S7.** A Pearson correlation r values between CSTB and NCS1 gene **A** regarding the autism class **B** regarding the control class.

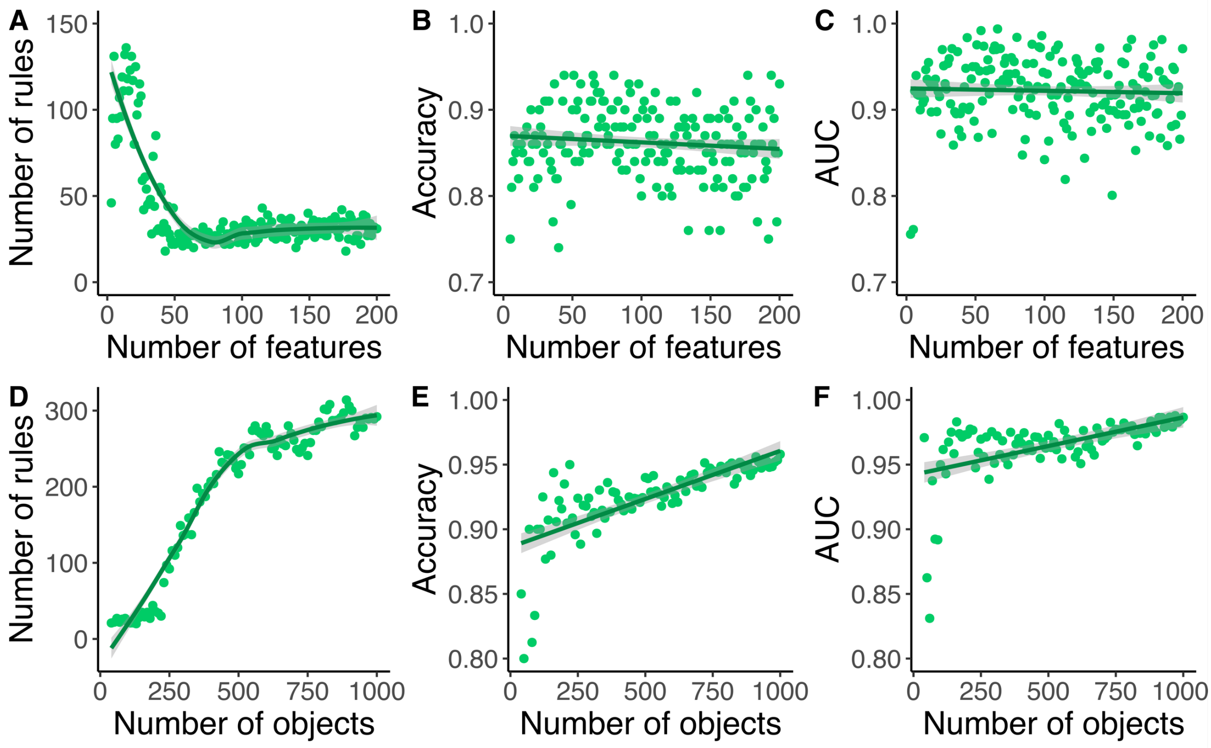

**Fig. S8.** The impact of features **A-C** and objects **D-F** number on the rule-based model quality. The tests were performed with the Johnson reduction algorithm on the synthetic data with a feature-feature correlation, rf = 0.4 and feature-decision correlation, rd = 0.6.

#### 3. Tables

**Table S1.** A comparison of the efficiency of the Johnson and Genetic reducers for basic (not recalculated) and recalculated rules. The data was discretized using the Equal Frequency method and the model was constructed with 10-fold CV of the standard voter classification method. The model was balanced using undersampling due to slightly imbalanced distribution of classes. The obtained p-values were Bonferroni-adjusted.

| Reducer type | Johnson | Genetic |
|---|---|---|
| Mean accuracy | 82% | 90% |
| Mean AUC | 0.85 | 0.98 |
| The total number of rules | 401 | 156650 |
| | basic | |
| Number of rules ns($p > 0.05$) | 123 | 156645 |
| Number of rules *($p \leq 0.05$) | 278 | 5 |
| Number of rules **$p \leq 0.01$ | 182 | 3 |
| Number of rules ***$p \leq 0.001$ | 104 | 2 |
| | recalculated | |
| Number of rules ns($p > 0.05$) | 218 | 156641 |
| Number of rules *($p \leq 0.05$) | 183 | 9 |
| Number of rules **($p \leq 0.01$) | 111 | 1 |
| Number of rules ***($p \leq 0.001$) | 39 | 0 |

**Table S2.** Performance evaluation for the Genetic reduction method. The average statistic values of support, accuracy and coverage are presented in the table. Top ranked co-predictors were selected as the most significant by Bonferroni-adjusted p-value.

| class | control | | autism | |
|---|---|---|---|---|
| total number of rules | 75530 | | 81120 | |
| rules | basic | recalculated | basic | recalculated |
| number of rules *($p \leq 0.05$) | 2 | 8 | 3 | 1 |
| LHS support | 4 | 6 | 5 | 6 |
| RHS support | 4 | 5 | 4 | 4 |
| accuracy | 0.81 | 0.80 | 0.86 | 0.74 |
| LHS coverage | 4% | 7% | 4% | 9% |
| RHS coverage | 6% | 6% | 7% | 7% |
| top ranked co-predictors | MAP7, NCKAP5L | MAP7, NCKAP5L | MAP7, COX2 | TCP11L1,CLDN17 RHPN1,PPOX |

**Table S3.** The performance evaluation of vote normalization methods in reclassifying the autism-control dataset. Bonferroni-adjusted p-value $\leq 0.05$ based filtration was investigated for the Johnson and Genetic reducers. The vote counts were normalized by different factors: median, mean, maximum (max), square root of the sum of squares (srss) or rule number (rulnum). The values represent accuracy (ACC) and AUC measures.

| method | none | | | | median | | | | mean | | | | max | | | | srss | | | | rulnum | | | |
|---|---|---|---|---|---|---|---|---|---|---|---|---|---|---|---|---|---|---|---|---|---|---|---|---|
| rules | basic | | recalculated | | basic | | recalculated | | basic | | recalculated | | basic | | recalculated | | basic | | recalculated | | basic | | recalculated | |
| quality | ACC | AUC | ACC | AUC | ACC | AUC | ACC | AUC | ACC | AUC | ACC | AUC | ACC | AUC | ACC | AUC | ACC | AUC | ACC | AUC | ACC | AUC | ACC | AUC |
| Johnson *($p \leq 0.05$) | 95% | 0.95 | 96% | 0.96 | 95% | 0.95 | 92% | 0.93 | 96% | 0.96 | 96% | 0.96 | 96% | 0.96 | 96% | 0.96 | 96% | 0.96 | 96% | 0.96 | 90% | 0.91 | 96% | 0.95 |
| Genetic *($p \leq 0.05$) | 71% | 0.73 | 49% | 0.55 | 71% | 0.71 | 67% | 0.70 | 72% | 0.73 | 67% | 0.70 | 72% | 0.73 | 67% | 0.70 | 72% | 0.73 | 67% | 0.70 | 70% | 0.70 | 49% | 0.55 |

**Table S4.** A comparison of the R.ROSETTA package to other R packages that enable rule-based classification modelling. The average accuracy, AUC, number of rules and time was calculated from the models with 20 repetitions of 10-fold CV without undersampling. The tests were performed on the autism-control dataset. The time was measured with tictoc library (Izrailev, 2014) as a time required for building a model to calculate the model quality measures.

| R package | C50 | RoughSets | | | R.ROSETTA | | RWeka | | |
|---|---|---|---|---|---|---|---|---|---|
| Package author | *Kuhn, M. et al. (2018)* | *Riza, L. S. et al. (2014)* | | | *Garbulowski, M. et al. (2020)* | | *Hornik, K., Buchta, C. and Zeileis, A. (2009)* | | |
| Algorithm abbreviation | C50 | AQ | CN2 | LEM2 | GenR | JohnR | JRip | OneR | PART |
| Detailed name of the algorithm | C5.0 | AQ | CN2 | Learning from Examples Module – version 2 | Genetic reducer | Johnson reducer | Repeated Incremental Pruning to Produce Error Reduction – RIPPER | 1R classifier | partial decision trees-based |
| Algorithm author | *Quinlan, J.R. (1992)* | *Michalski, R.S. et al. (1991)* | *Clark, P.E. and Niblett, T. (1989)* | *Grzymala-Busse J.W. (1997)* | *Johnson, D.S. (1974)* | *Wroblewski, J. (1995)* | *Cohen, W.W. (1995)* | *Holte, R.C. (1993)* | *Frank, E. and Witten, I.H. (1998)* |
| Function | C5.0.default | AQRules.RST | RI.CN2Rules.RST | RI.LEM2Rules.RST | rosetta | | Jrip | OneR | PART |
| Discretization | equal frequency | | | | | | | | |
| accuracy | 75% | 75% | 76% | 76% | 91% | 82% | 72% | 62% | 75% |
| AUC | 0.74 | 0.74 | 76% | 0.74 | 0.99 | 0.88 | 0.72 | 0.61 | 0.75 |
| number of rules | 10 | 40 | 14 | 19 | 76974 | 185 | 5 | 1 | 9 |
| time[s] | 0,8 | 8,3 | 13,1 | 1,9 | 1529,8 | 3,9 | 1,3 | 1,1 | 1,2 |

**Table S5.** A correlation between the outcome and clinical data of the autism-control dataset. The p-values were estimated with the Student's t-test. Non-significant differences are shown between batches and parental ages. The highly significant correlation for the age of subjects and outcome is shown.

| Class | control | autism | p-value |
|---|---|---|---|
| Number of the samples | 64 | 82 | - |
| Subject age | 7.9±2.1 | 5.5±2.1 | $4.26\times10^{-9}$ |
| Maternal age | 29.7±5 | 30.1±5.6 | 0.71 |
| Paternal age | 30.3±5.3 | 31.5±6.3 | 0.27 |
| Batch | B1(32), B2(32) | B1(36), B2(46) | 0.46 |

**Table S6.** A comparison of the performance of four dimensionality reduction methods applied to the autism-control dataset. For Boruta and Student's t-test the p-value ≤ 0.05 threshold was established. The FCBF method selected genes with the Information Gain (IG) greater than 0. Monte Carlo Feature Selection (MCFS) estimated a Relative Importance (RI) threshold with a critical angle method.

| Feature selection method | Boruta | FCBF | MCFS | Student's t-test |
|---|---|---|---|---|
| R package | Boruta | Biocomb | rmcfs | stats |
| Threshold | $p < 0.05$ | IG > 0 | RI > 0.036 | $p < 0.05$ |
| Number of features | 12 | 35 | 16 | 13 |
| Discretization | No | Yes | No | No |

**Table S7.** The result of the FCBF method applied on the autism-control dataset. The list is decreasingly sorted from the features with the highest IG. The position of each feature in a ranking is given in the first column. The translation between microarray probe ID and gene ID is shown in the last two columns.

| position | IG | gene ID | probe ID |
|---|---|---|---|
| 1 | 0,10860441 | MAP7 | 202890_at |
| 2 | 0,0995046 | COX2 | 1553569_at |
| 3 | 0,094478 | NCKAP5L | 1562457_at |
| 4 | 0,08567925 | ZSCAN18 | 217593_at |
| 5 | 0,08391652 | RHPN1 | 235998_at |
| 6 | 0,07773368 | PPOX | 238118_s_at |
| 7 | 0,07625543 | NPR2 | 204310_s_at |
| 8 | 0,07608236 | NCS1 | 222570_at |
| 9 | 0,07128859 | PSMG4 | 233443_at |
| 10 | 0,06793192 | SCIN | 1552367_a_at |
| 11 | 0,06564102 | CSTB | 236449_at |
| 12 | 0,06244449 | TSPOAP1 | 205839_s_at |
| 13 | 0,06222545 | TCP11L1 | 205796_at |
| 14 | 0,06185506 | 234817_at | 234817_at |
| 15 | 0,06116627 | TMLHE-AS1 | 1560797_s_at |
| 16 | 0,0606206 | PSMD4 | 200882_s_at |
| 17 | 0,05976664 | ZFP36L2 | 201367_s_at |
| 18 | 0,05938284 | B3GNT7 | 1555962_at |
| 19 | 0,05752864 | MSI2 | 225238_at |
| 20 | 0,05732106 | CAPS2 | 224370_s_at |
| 21 | 0,05700458 | MIR646HG | 1562051_at |
| 22 | 0,05581372 | CLDN17 | 221328_at |
| 23 | 0,05510055 | BAHD1 | 203051_at |
| 24 | 0,05288485 | OR51B5 | 1570516_s_at |
| 25 | 0,050778 | C11orf95 | 218641_at |
| 26 | 0,0490634 | ATXN8OS | 216404_at |
| 27 | 0,04738852 | NRG2 | 242303_at |
| 28 | 0,04694185 | LOC400655 | 216703_at |
| 29 | 0,04694185 | GJA9 | 221415_s_at |
| 30 | 0,0446669 | VPS8 | 234028_at |
| 31 | 0,04336461 | FLRT2 | 240259_at |
| 32 | 0,0388937 | C1QTNF7 | 239349_at |
| 33 | 0,03606374 | KLF8 | 219930_at |
| 34 | 0,03568021 | CWF19L2 | 1566515_at |
| 35 | 0,03116567 | DEPDC1 | 222958_s_at |
